# Identification of bis-benzylisoquinoline alkaloids as SARS-CoV-2 entry inhibitors from a library of natural products in vitro

**DOI:** 10.1101/2020.12.13.420406

**Authors:** Chang-Long He, Lu-Yi Huang, Kai Wang, Chen-Jian Gu, Jie Hu, Gui-Ji Zhang, Wei Xu, You-Hua Xie, Ni Tang, Ai-Long Huang

## Abstract

Coronavirus disease 2019 (COVID-19) caused by severe acute respiratory syndrome coronavirus 2 (SARS-CoV-2) is a major public health issue. To screen for antiviral drugs for COVID-19 treatment, we constructed a SARS-CoV-2 spike (S) pseudovirus system using an HIV-1-based lentiviral vector with a luciferase reporter gene to screen 188 small potential antiviral compounds. Using this system, we identified nine compounds, specifically, bis-benzylisoquinoline alkaloids, that potently inhibited SARS-CoV-2 pseudovirus entry, with EC_50_ values of 0.1–10 μM. Mechanistic studies showed that these compounds, reported as calcium channel blockers (CCBs), inhibited Ca^2+^-mediated membrane fusion and consequently suppressed coronavirus entry. These candidate drugs showed broad-spectrum efficacy against the entry of several coronavirus pseudotypes (SARS-CoV, MERS-CoV, SARS-CoV-2 [S-D614, S-G614, N501Y.V1 and N501Y.V2]) in different cell lines (293T, Calu-3, and A549). Antiviral tests using native SARS-CoV-2 in Vero E6 cells confirmed that four of the drugs (SC9/cepharanthine, SC161/hernandezine, SC171, and SC185/neferine) reduced cytopathic effect and supernatant viral RNA load. Among them, cepharanthine showed the strongest anti-SARS-CoV-2 activity. Collectively, this study offers new lead compounds for coronavirus antiviral drug discovery.

## Introduction

Coronavirus disease 2019 (COVID-19) caused by severe acute respiratory syndrome coronavirus 2 (SARS-CoV-2) is a major public health issue. The spike (S) protein mutation D614G (a single change in the genetic code; D = aspartic acid, G = glycine) became dominant in SARS-CoV-2 during global pandemic, which displayed increased infectivity.^1^ Entry of a virus into host cells is one of the most critical steps in the viral life cycle. Since blockade of the entry process is a promising therapeutic option for COVID-19, research attention has been focused on the discovery of viral entry inhibitors. Although SARS-CoV-2 entry inhibitor development is very attractive, no candidates have progressed into clinical trials yet.

## Materials and Methods

### Plasmids

The codon-optimized gene encoding SARS-CoV-2 spike (S) protein (GenBank: QHD43416) with 19 amino acids deletion at the C-terminal was synthesized by Sino Biological Inc (Beijing, China), and cloned it into the pCMV3 vector between the restriction enzyme *Kpn*I and *Xba*I sites (denoted as pS-D614). The recombinant plasmid pS-D614 was used as template, and the D614G mutant S-expressing plasmid (denoted as pS-G614) was constructed by site-directed mutagenesis. N501Y.V1 (Variant 1) mutant Spike proteins of SARS-CoV-2 were codon-optimized and synthesized by GenScript Inc (Nanjing, China) and cloned into pCMV3 vector (denoted as pS-Variant1). N501Y.V2 (Variant 2) mutant Spike-expressing plasmid (denoted as pS-Variant2) was constructed by site-directed mutagenesis, with pS-D614 plasmid as a template. SARS-CoV S-expressing plasmid (Cat: VG40150-ACGLN, named as pS-SARS) and MERS-CoV S-expressing plasmid (Cat: VG40069-CF, named as pS-MERS) were obtained from Sino Biological Inc (Beijing, China). The VSV-G-expressing plasmid pMD2.G was donated by Prof. Ding Xue from Tsinghua University (Beijing, China). The HIV-1 NL4-3 ΔEnv Vpr luciferase reporter vector (pNL4-3.Luc.R-E-) constructed by N. Landau was donated by Prof. Cheguo Cai from Wuhan University (Wuhan, China). Human ACE2 and DPP4 expression plasmids were derived from GeneCopoeia (Guangzhou, China).

### Cell lines and cell culture

HEK 293T, A549, and Calu3 cells were purchased from the American Type Culture Collection (ATCC, Manassas, VA, USA). Cells were cultured at 37 °C and 5% CO_2_ atmosphere in Dulbecco’s modified Eagle medium (DMEM; Hyclone, Waltham, MA, USA) containing 10% fetal bovine serum (FBS; Gibco, Rockville, MD, USA), 100 mg/mL of streptomycin, and 100 units/mL of penicillin. HEK 293T cells transfected with human ACE2 and DPP4 (293T-ACE2, 293T-DPP4) were cultured under the same conditions with the addition of G418 (0.5 mg/mL) to the medium.

### Antigens and antibodies

The RBD domain of SARS-CoV-2 S protein (His-tag) were synthesized by Prof. Xuefei Cai at Key Laboratory of Molecular Biology for Infectious Diseases (Ministry of Education), Chongqing Medical University. The anti-RBD monoclonal antibody was kindly provided by Prof. Aishun Jin from Chongqing Medical University. RBD-binding peptide SBP1 derived from human ACE2 α helix 1 (Ac-IEEQAKTFLDKFNHEAEDLFYQS-NH_2_) was synthesized by GenScript (Nanjing, China).

### Compounds and reagents

Custom compound library containing 188 small molecules, remdesivir, and aloxistatin (E-64d) were purchased from MedChemExpress (HY-L027) and Chemdiv. BAPTA-AM (T6245) was purchased from TargetMol. All the compounds were dissolved in dimethyl sulfoxide (DMSO) at a stock concentration of 20 mM.

### Cell cytotoxicity assay

The CellTiter 96® AQ_ueous_ One Solution Cell Proliferation Assay (G3582, Promega, USA) was used to assess cell viability according to the product’s description. Briefly, HEK 293T cells were dispensed into 96-well plate (2×10^4^ cells/well), cultured in medium containing gradient concentrations of the compound for 72 hours at 37 °C in a humidified 5% CO_2_ incubator. The cells were incubated with 100 µl fresh medium after removal of the medium. Then 20 µl of CellTiter 96® AQueous One Solution Reagent was added into each sample well and incubating the plate was incubated at 37°C for 1-4 hours in a humidified 5% CO_2_ atmosphere. The absorbance at 490 nm was measured using a microplate reader (Synergy H1, BioTek, USA).

### Pseudovirus production and quantification

5×10^6^ HEK 293T cells were co-transfected with 6 μg each of pNL4-3.Luc.R-E-and recombinant plasmid (pS-SARS, pS-MERS, pS-D614, pS-G614, pS-Variant1, or pS-Variant2) using Lipofectamine 3000 Transfection Reagent (Invitrogen, Rockville, MD) according to the manufacturer’s instructions. After 48 h transfection, pseudotyped viruses expressing S-SARS, S-MERS, S-D614, S-G614, N501Y.V1, and N501Y.V2 spike protein were harvested, centrifuged and filtered through 0.45 μm filters, and subsequently stored at -80°C. 293T cells were co-transfected with pNL4-3.Luc.R-E- and pMD2.G plasmid to collect the VSV-G pseudovirus.

The copies of the pseudovirus were expressed as numbers of viral RNA genomes per mL of viral stock solution and determined using RT-qPCR with primers and a probe that targeting LTR. Sense primer: 5′-TGTGTGCCCGTCTGTTGTGT-3′, anti-sense primer: 5′-GAGTCCTGCGTCGAGAGAGC-3′, probe: 5′-FAM-CAGTGGCGCCCGAACAGGGA-BHQ1-3′. Briefly, viral RNAs were extracted according to the manufacturer’s instructions with TRIzol reagent (Invitrogen). Then, the TaqMan One-Step RT-PCR Master Mix Reagents (Applied Biosystems, Thermo Fisher) was used to amplify total RNAs. pNL4-3.Luc.R-E-vector with certain copies was used to generate standard curves. All the pseudotyped viruses were titrated to the uniform titer (copies/mL) for the following research.

### Compound screening

For pseudovirus-based inhibition assay, 188 compounds were screened via luciferase activity. HEK 293T cells cultured in 96-well plate (2×10^4^ cells/well) were incubated with each compound (20 μM) for 1 hour and were infected with the same amount of pseudovirus (3.8 × 10^4^ copies in 50 μL). The cells were replaced to fresh DMEM medium 8 h post-infection. Cells were lysed by 30 μl lysis buffer (Promega, Madison, WI, USA) at 72 h post-infection to measure RLU with luciferase assay reagent (Promega, Madison, WI, USA) according to the product description. All data were performed at least three times and expressed as means ± standard deviations (SDs).

### Cell-cell fusion assay

HEK 293T effector cells were transfected with plasmid pAdTrack-TO4-GFP encoding green fluorescent protein (GFP) or pS-G614 encoding the corresponding SARS-CoV-2 S protein. 293T-ACE2 cells were used as target cells. At 8 h post-transfection, the effector cells were washed twice with PBS and were pretreated with compounds or DMSO as control for another 16h. Subsequently, the effector cells were overlaid on target cells at a ratio of approximately one S-expressing cell to two receptor-expressing cells with about 90% confluent. After a 4-hour coculture, images of syncytia were captured with an inverted fluorescence microscope (Nikon eclipse Ti, Melville, NY).

For quantification of cell-cell fusion, three fields were randomly selected in each well to count the fused and unfused cells. The fused cells were at least twice as large as the unfused cells, and the fluorescence intensity was weaker in fused cells since GFP diffusion from one effector cell to target cells. The percentage of cell-cell fusion was calculated as: [(number of the fused cells/number of the total cells) × 100%].

### Competitive ELISA

The recombinant RBD proteins derived from SARS-CoV-2 were coated on 96-well microtiter plate (50ng/well) at 4°C overnight. After blocked with blocking buffer (5% FBS and 2% BSA in PBS) for 1 hour at 37°C, serial dilution solutions of compounds, ACE2 peptide SBP1 or DMSO were added into the plates and incubated at 37°C for 1 hour. Plates were washed five times with phosphate-buffered saline, 0.05% Tween-20 (PBST) to remove the free drug or DMSO. The wells were incubated with mouse anti-RBD monoclonal antibody (1:1000 dilution) for 1 hour at 37°C, and then washed with PBST five times and incubated with Horseradish peroxidase (HRP)-conjugated goat anti-mouse antibody (Abmart, Shanghai, China) for 1 hour at 37°C. TMB substrate was added and incubated for 15 minutes at 37°C for color development, finally the absorbance at 450 nm was measured by a microplate reader.

### Differential scanning fluorimetry

Briefly, purified His-RBD was diluted to 200 μg/mL in PBS buffer containing Sypro Orange 5× (ThermoFisher) in a 96-well white PCR plate. Compounds and ACE2 derived peptide SBP1 were added at a final concentration of 20 μM. All samples were tested in triplicate with a Bio-Rad RT-PCR system. The samples were first equilibrated at 25 °C for 3 min, then heated from 25 °C to 85 °C with a step of 1 °C per 1 min, and the fluorescence signals were continuously collected using CFX Maestro.

### Functional analysis of Ca^2+^ in pseudovirus infectivity

293T-ACE2 cells were seeded in 96-well plate (2×10^4^ cells/well) and incubated at 37 °C for 8h. For extracellular calcium depletion assays, cells were washed three times using PBS with or without Ca^2+^. Calcium-free medium with or without 2 mM calcium chloride and 5 μM compounds were added and incubated for 1h at 37°C. Next, S-G614 pseudovirus was added to the cells for 8 h at 37°C. For intracellular calcium depletion assays, cells were washed three times using PBS with or without Ca^2+^, DMEM with BAPTA-AM (20 μM) or DMSO and gradient concentrations of compounds were added and incubated for 2 h at 37°C. In the following, S-G614 pseudovirus was added to infect the cells for 8 h at 37°C. For both types of assays, complete medium was then added after removed the medium. The cells were measured by luciferase activity.

### Cellular cholesterol measurement

Cellular cholesterol was measured using the Cholesterol/Cholesterol Ester-Glo™ Assay (J3190, Promega, USA) according to the product’s description. In short, HEK 293T-ACE2 cells were assigned to 96-well plates (4×10^4^ cells/well) and cultured for 24 hours at 37 °C with 5 μM compounds. DMSO was used as negative control. The medium in the 96-well plate was removed and cells was washed twice with 200 µl PBS. Then, 50 µl of cholesterol lysis solution was added, shaking the plate carefully and incubating for 30 minutes at 37°C. Following, 50 µl of cholesterol detection reagent was added with esterase or without esterase to all wells and the plate was incubated at room temperature for 1 hour. Finally, recording luminescence with a plate-reading luminometer. Total cholesterol concentration was calculated by comparing the luminescence of samples and controls under the same condition.

### Cytopathic effect (CPE) assay and quantification of SARS-CoV-2 infection

Vero E6 cells were dispensed into 96-well plate (4.0×10^4^ cells/well), pre-treated with medium containing compounds or DMSO for 1 hour at 37 °C. Then 60 μl DMEM medium which supplemented with compounds or DMSO was replaced immediately, the same amount of SARS-CoV-2 (100 TCID_50_/well) was added and incubated for 1 hour at 37 °C. The mixture was removed and the cells were washed twice with PBS, and cultured with 100 μl fresh medium for 48 hours at 37 °C with 5% CO_2_ atmosphere. Cytopathic effects (CPE) induced by the virus was observed using microscope at 48 h post-inoculation. Cell culture supernatant was collected at 48 h post-infection for viral RNA quantification using the novel coronavirus real-time RT-PCR Kit (Shanghai ZJ Bio-Tech Co, Ltd, Shanghai, China). Remdesivir (5 μM) was used as positive control in the experiment.

### Statistical analyses

Data were analyzed using GraphPad Prism version 6.0 software and were presented as means ± SD. Statistical significance was determined using ANOVA for multiple comparisons. Student’s t test was applied to compare the two groups. Differences with P values < 0.05 were deemed statistically significant.

## Results

Using a luciferase-expressing pseudovirus encoding SARS-CoV-2 S (G614) protein, a library of 188 natural compounds (Supplementary Tab. S1) was screened in 293T-ACE2 cells (HEK 293T cells overexpressing human angiotensin-converting enzyme 2) to find novel anti-SARS-CoV-2 entry inhibitors. A workflow chart of screening is shown in Fig. 1a. After a preliminary screening, 41 compounds associated with a relative infection rate <30% (Fig. 1b) were identified. We selected 19 compounds with low cytotoxicity for further testing (Supplementary Tab. S2, Fig. S1). Among the 19 hits, nine compounds (SC9, SC161, SC171, SC182–187) with relatively high activity (EC_50_ < 10 μM), low cytotoxicity (CC_50_ > 20 μM), and high specificity (SI > 10) were selected for subsequent analyses. Specifically, all these compounds were bis-benzylisoquinoline alkaloids except SC171.

**Fig 1.**
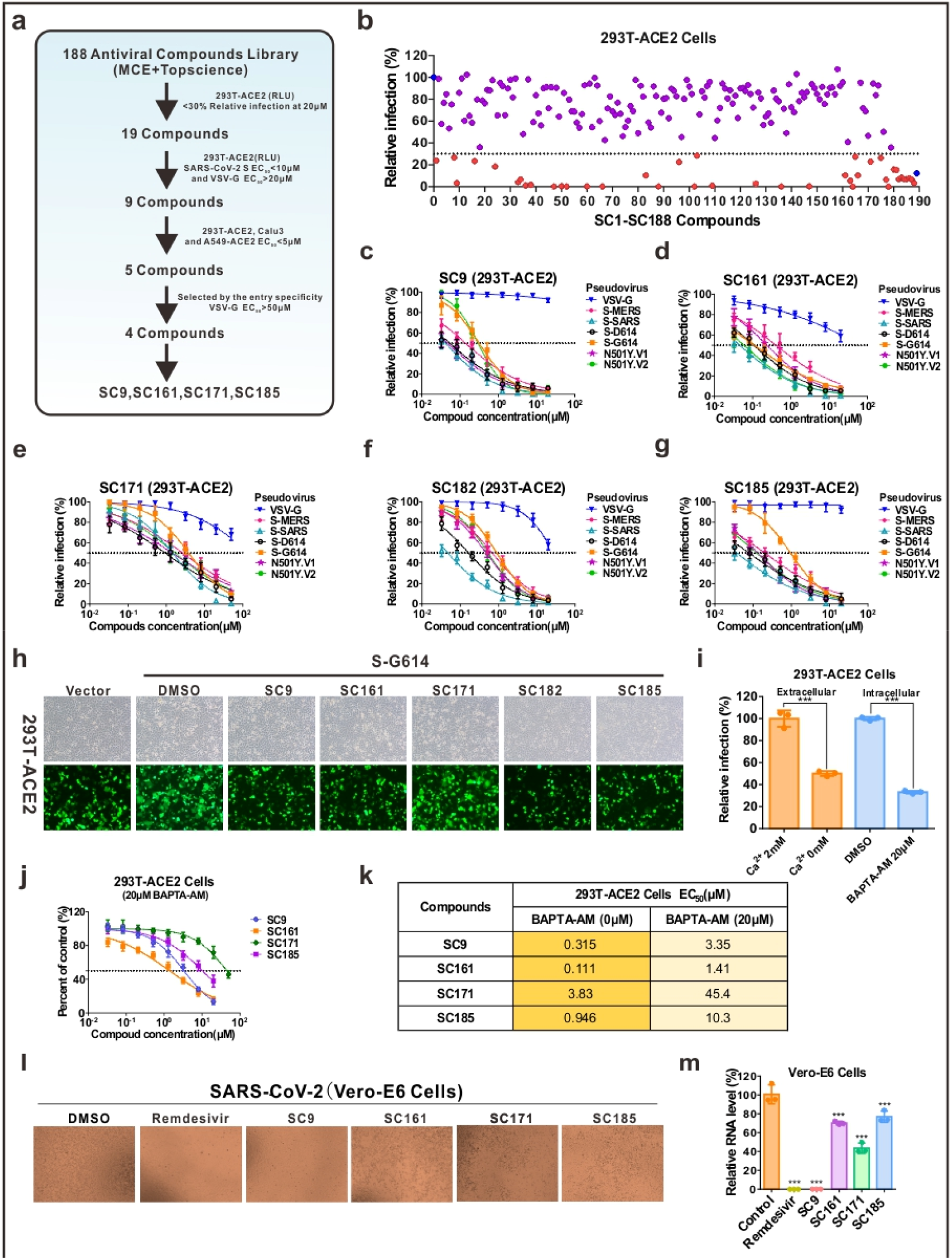
Identification of bis-benzylisoquinoline alkaloids as SARS-CoV-2 entry inhibitors. **a** Schematic diagram of the screening workflow with selection criteria for hits outlined. **b** Scatter plot of primary screening of 188 compounds against S-G614 infection. Inhibition ratios for all drugs obtained in a preliminary screening are represented by scattered points. Red dots indicate the 41 compounds with an inhibition rate ≥70%. DMSO and aloxistatin (blue dot) were used as a negative and positive control, respectively. **c-g** Dose-response curves of five selected compounds (**c**) SC9, (**d**) SC161, (**e**) SC171, (**f**) SC182, (**g**) SC185 on VSV-G, S-D614, S-G614, S-SARS, S-MERS, N501Y.V1, and N501Y.V2 pseudoviruses. **h** Inhibitory effect of SC9, SC161, SC171, SC182, and SC185 at 5 μM on SARS-CoV-2 S mediated cell-cell fusion. **i** Effect of extracellular and intracellular Ca^2+^ depletion on S-G614 pseudovirus entry in 293T cells. **j-k** Inhibition curves (**j**) and EC_50_ values (**k**) of the compounds against S-G614 pseudovirus entry in the presence of 20 μM BAPTA-AM. **l** The inhibitory effect of the compounds on native SARS-CoV-2 infection by observing their cytopathogenic effects. SC9, SC161, SC171, and SC185 were tested at 10 μM, and DMSO and remdesivir (5 μM) were used as a negative and positive control, respectively. **m** The relative viral RNA levels in the SC9, SC161, SC171, and SC185 (10 μM) treatment groups were 0.08%, 70.27%, 43.55%, and 76.98% respectively. \**P* < 0.05; \*\**P* < 0.01; \*\*\**P* < 0.001. All experiments were repeated at least three times.

Next, we analyzed the relationship between the antiviral efficacy of the nine selected compounds against S-G614 pseudovirus and the timing of treatment (Supplementary Fig. S2). We divided the S-G614 pseudovirus entry into three stages: pretreatment, pseudovirus entry, and pseudovirus post-entry. In total, eight experimental groups were set up for each compound, including seven treatment groups (A–G) and a control group. Importantly, pretreatment with each compound (group B) significantly inhibited S-G614 pseudovirus infection. In the pseudovirus entry stage (group C), the compounds exerted similar suppressive effects. However, in the pseudovirus post-entry stage (group D), none of the compounds showed any inhibitory effect. Our data demonstrated that the nine selected compounds showed effective entry blockade presenting in the pretreatment stage, indicating that they target host factors at the entry stage.

Cell lines mimicking important aspects of respiratory epithelial cells should be used when analyzing the anti-SARS-CoV-2 activity. Hence, we determined their EC_50_ values against S-G614 pseudovirus in Calu-3 and A549 cells (Supplementary Fig. S3a-i). Five compounds (SC9, SC161, SC171, SC182, and SC185) with EC_50_ < 10 μM in all three cells lines were selected for subsequent experiments.

To determine whether these compounds have broad-spectrum antiviral effects against coronaviruses, we constructed S-D614, N501Y.V1 (B.1.1.7), N501Y.V2 (B. 1.351), S-SARS, and S-MERS pseudoviruses using the same lentiviral system as S-G614, and then determined the EC_50_ values of SC9 (cepharanthine, Fig. 1c), SC161 (hernandezine, Fig. 1d), SC171 (Fig. 1e), SC182 (tetrandrine, Fig. 1f), and SC185 (neferine, Fig. 1g) against these pseudoviruses in 293T cells expressing ACE2 or dipeptidyl peptidase 4 (DPP4). Interestingly, SC9, SC161, SC171, and SC185 exhibited highly potent pan-inhibitory activity against S-pseudotyped coronaviruses including two emerging SARS-CoV-2 variants N501Y.V1 and N501Y.V2, reported in United Kingdom and South Africa (Supplementary Fig. S3j). As SARS-CoV and SARS-CoV-2 have been reported to enter host cells via binding to ACE2, and while DPP4 is critical for MERS-CoV entry, it could be ruled out that the five compounds interfere with ACE2 to block pseudovirus entry.

Then, we used competitive ELISAs and thermal shift assays to determine whether these five compounds interact with the receptor-binding domain (RBD) in S protein of SARS-CoV-2. SBP1, a peptide derived from the ACE2 α1 helix, bound RBD of SARS-CoV-2 and exhibited a weak ability to inhibit the entry of S-G614 pseudovirus (Supplementary Fig. S4a), whereas the interaction between SC9, SC161, SC171, or SC185 and RBD was negligible (Supplementary Fig. S4c-d). Thus, the blockade of virus entry by these candidate compounds is not related to interaction with the SARS-CoV-2 RBD domain.

Following attachment to host receptor, the membrane fusion process mediated by S protein of SARS-CoV-2 plays an important role in viral entry. Thus, we examined whether these compounds perturb cell fusion mediated by SARS-CoV-2 S protein. Compared to DMSO, SC9, SC161, SC182, and SC185 at 5 μM potently inhibited SARS-CoV-2 S-mediated membrane fusion of 293T cells with approximately 90% decrease of fusion rates (Fig. 1h, Supplementary Fig. S4e). Previous studies have shown that the calcium ion (Ca^2+^) plays a critical role in SARS-CoV or MERS-CoV S-mediated fusion with host cells.^2^ Calcium channel blockers (CCBs), originally used to treat cardiovascular diseases, are supposed to have a high potential to treat SARS-CoV-2 infections.^3^ We noted that the identified bis-benzylisoquinoline alkaloids had been reported as CCBs.^4^ Herein, the bis-benzylisoquinoline CCBs abolished S–ACE2-mediated membrane fusion in 293T-ACE2 cells. Calcium-free medium or intracellular Ca^2+^ chelation with BAPTA-AM significantly diminished SARS-CoV-2 pseudovirus infection (Fig. 1i, Supplementary Fig. S4f-i), suggesting that Ca^2+^ is required for SARS-CoV-2 entry. Upon pretreatment with BAPTA-AM, the bis-benzylisoquinoline CCBs had approximately 10-fold higher EC_50_ values than those without BAPTA-AM pretreatment (Fig. 1j-k, Supplementary Fig. S4j-k). Additionally, perturbation of the cholesterol biosynthesis pathway with the CCB amlodipine reduced viral infection.^5^ Consistent herewith, the bis-benzylisoquinoline CCBs upregulated intracellular cholesterol (Supplementary Fig. S4i), which also likely contributes to inhibition of viral infection. These data indicated that blockade of S-G614 pseudovirus entry by bis-benzylisoquinoline CCBs mainly depends on calcium homeostasis.

Finally, the antiviral activities of SC9 (cepharanthine), SC161 (hernandezine), SC171, and SC185 (neferine) were confirmed in Vero E6 cells infected with native SARS-CoV-2. SARS-CoV-2 induced cytopathogenic effect (CPE) and the viral RNA levels were partly inhibited by these compounds, with SC9 (cepharanthine) at the highest efficacy and the other three compounds much less active. (Fig. 1l-m). The results showed that these compounds inhibited SARS-CoV-2 to varying degrees and may be useful as leads for SARS-CoV-2 therapeutic drug development.

## Discussion

In summary, we reported a set of bis-benzylisoquinoline alkaloids as coronavirus entry inhibitors. These inhibitors effectively protected different cell lines (293T, Calu-3, and A549) from infection by different coronaviruses (SARS-CoV, MERS-CoV, SARS-CoV-2 [S-D614 and S-G614 variant]) *in vitro*. The compounds block host calcium channels, thus inhibiting Ca^2+^-mediated fusion and suppressing virus entry. Considering the effectiveness of CCBs in the control of hypertension, our study provided clues to support that CCBs may be useful in treating coronavirus infection in patients with hypertension.

## Acknowledgments

We would like to thank Prof. Cheguo Cai (Wuhan University, Wuhan, China) for providing the pNL4-3.Luc.R-E-plasmid. This work was supported by the Key Laboratory of Infectious Diseases, CQMU, 202001, the Science and Technology Research Program of Chongqing Municipal Education Commission (KJQN202000418 to L-Y.H.), Emergency Project from the Science & Technology Commission of Chongqing (cstc2020jscx-fyzx0053, cstc2020jscx-dxwtB0050 to A-L.H.), and the Emergency Project for Novel Coronavirus Pneumonia from the Chongqing Medical University (CQMUNCP0302 to K.W.).

## Author contributions

A.L.H., N.T., and Y.H.X. designed and directed the study. C.L.H., L.Y.H., K.W., J.H., and G.J.Z. constructed the pseudoviruses and screened the compounds. C.J.G. and W.X. performed authentic SARS-CoV-2 assays. All authors reviewed the manuscript and consented to the description of author contribution.

## Additional information

The online version of this article contains supplementary material, which is available to authorized users.

## Competing interests

The authors declare no competing interests.

## Notes

### Competing Interest Statement

The authors have declared no competing interest.

### Summary of Updates

S-pseudotyped coronaviruses including two emerging SARS-CoV-2 variants N501Y.V1 and N501Y.V2, reported in United Kingdom and South Africa, were tested. Figure 1 revised; Figures S1 to S4 revised;Table S1 to S2 updated.

