## Supplementary for "Identification of bis-benzylisoquinoline alkaloids as SARS-CoV-2 entry inhibitors from a library of natural products in vitro"

**This PDF file includes:**

Figures S1 to S4

Table S1 to S2

**Figure S1**

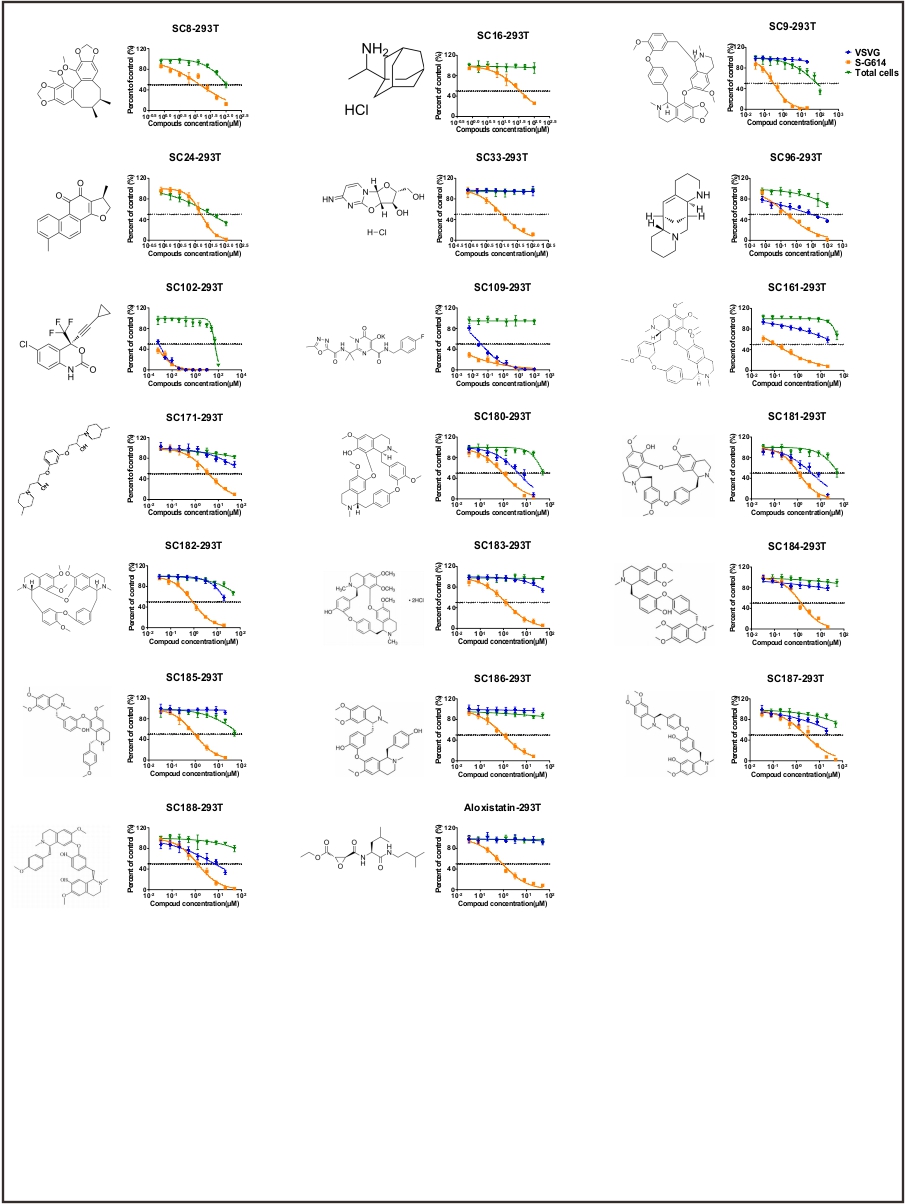

**Fig. S1.** The chemical structures and inhibitory activities of 19 hits and aloxistatin against SARS-CoV-2 S-G614, VSV-G pseudoviruses and cell viability assay of 293T-ACE2.

**Figure S2**

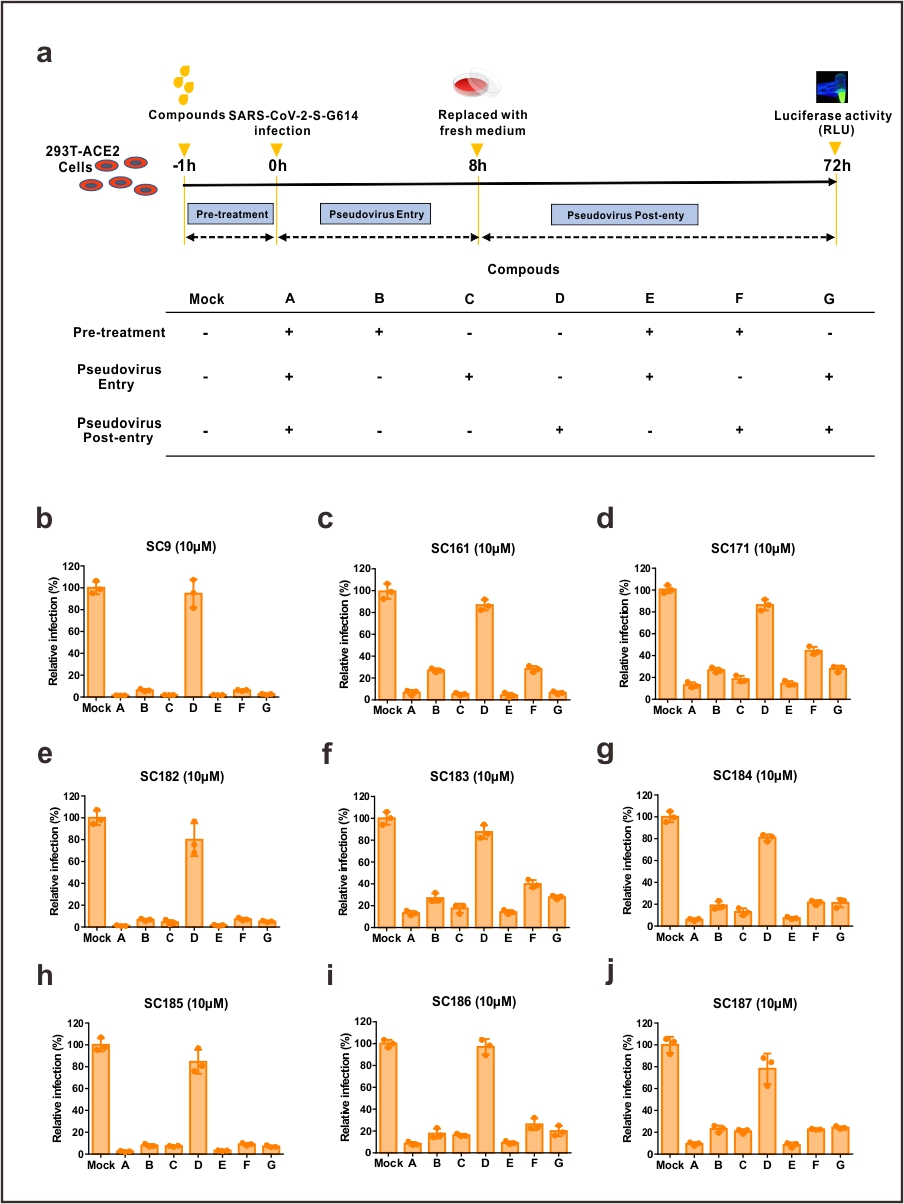

**Fig. S2** Effect of treatment timing on the efficacy of nine selected compounds on SARS-CoV-2 S-G614 pseudovirus entry. **a** Treatment timing diagrams for the nine selected compounds. HEK 293T cells were treated with the nine compounds or DMSO before, during, or after S-G614 pseudovirus entry. Seven treatment conditions (A–G) were tested for each compound. **b–j** Inhibitory effects of **(b)** SC9, **(c)** SC161, **(d)** SC171, **(e)** SC182, **(f)** SC183, **(g)** SC184, **(h)** SC185, **(i)** SC186, and **(j)** SC187 on the entry of S-G614 pseudovirus at different treatment time points. All experiments were repeated at least three times.

**Figure S3**

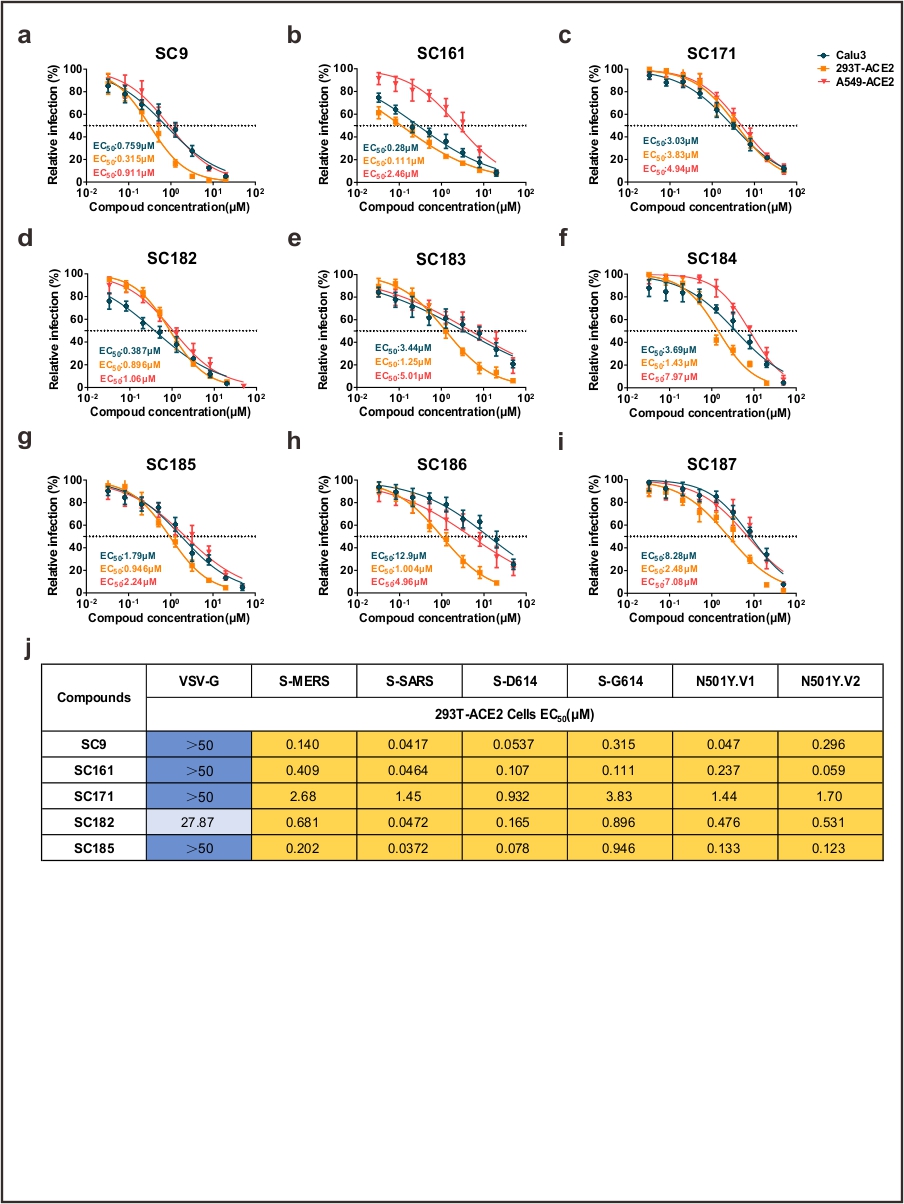

**Fig. S3** Efficacy of compounds in Calu3, 293T-ACE2, and A549 cell lines against different coronaviruses. **a-i** Inhibitory effects of **(a)** SC9, **(b)** SC161, **(c)** SC171, **(d)** SC182, **(e)** SC183, **(f)** SC184, **(g)** SC185, **(h)** SC186, and **(i)** SC187 against S-G614 infection in the three cell lines. All experiments were repeated at least three times. (**j**) EC_50_ values of five selected compounds against entry of different pseudoviruses. All experiments were repeated at least three times.

**Figure S4**

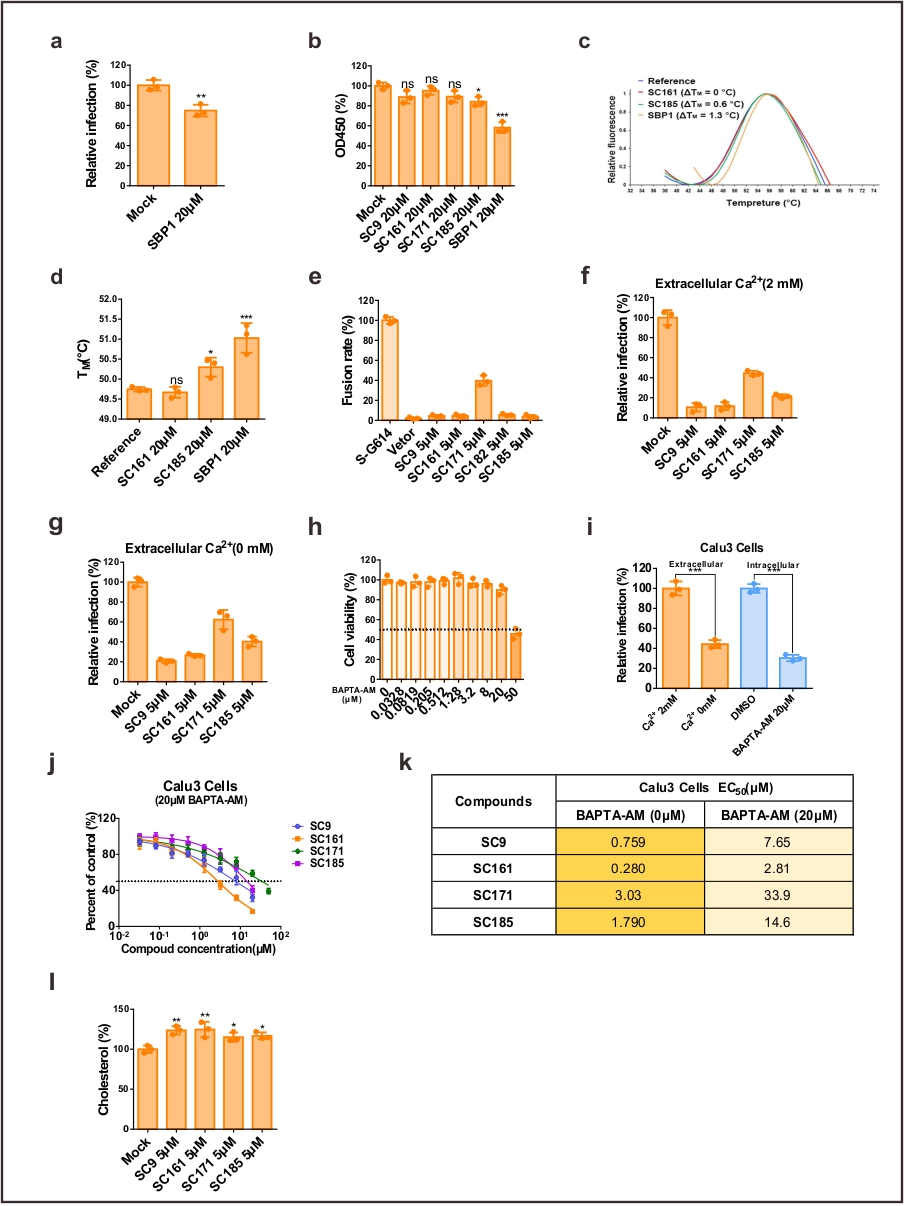

**Fig. S4** Investigation of the mechanism of the selected compounds. **a** Inhibitory effect of the ACE2 peptide SBP1 at 20 μM on the invasion of S-G614 pseudovirus. **b** Binding of pre-coated His-RBD with compounds at 20 μM as determined by competitive ELISA. SBP1 and DMSO were used as a positive and negative control, respectively. **c** Melting curves for His-RBD (10 μM) incubated with 20 μM of each compound tested by differential scanning fluorimetry. **d** T_M_ values of His-RBD incubated with SC161, SC185, or SBP1. **e** Cell–cell fusion rate in the presence of 5 μM SC9, SC161, SC171, SC182, or SC185. The fusion rate in a DMSO-treated group was set as 100%. **f** Effect of 5 μM of each compound on S-G614 pseudovirus entry into 293T-ACE2 cells in medium supplemented with 2 mM CaCl_2_. **g** Effect of 5 μM of each compound on S-G614 pseudovirus entry in calcium-depleted culture medium. **h** Cell viability assay of 293T-ACE2 cells treated with increasing concentrations of BAPTA-AM (0–50 μM). **i** Effect of extracellular and intracellular Ca^2+^ depletion on S-G614 pseudovirus entry in Calu-3 cells. **j-k** Inhibition curves (**j**) and EC_50_ values (**k**) of the compounds against S-G614 pseudovirus entry in the presence of 20 μM BAPTA-AM. **l** Cholesterol (normalized to total protein) in 293T-ACE2 cells treated with the compounds or DMSO for 24 h. **P* < 0.05; ***P* < 0.01; ****P* < 0.001. ns, not significant. All experiments were repeated at least three times.

**Table S1**

| **Supplementary Tab. 1** List of 188 compounds. | | | | | | | | |
| --- | --- | --- | --- | --- | --- | --- | --- | --- |
| **Code** | **Compounds Name** | **M. Wt** | **Code** | **Compounds Name** | **M. Wt** | **Code** | **Compounds Name** | **M. Wt** |
| SC1 | Xanthohumol | 354.40 | SC64 | 2,2'-Anhydrouridine | 226.19 | SC127 | 3,4-Dimethoxycinnamic acid | 208.21 |
| SC2 | Lycorine | 287.31 | SC65 | Tricin | 330.29 | SC128 | Fumagillin | 458.54 |
| SC3 | Oxindole | 133.15 | SC66 | Tubercidin | 266.25 | SC129 | Geniposide | 388.37 |
| SC4 | L-Lysine | 146.19 | SC67 | (-)-Epicatechin gallate | 442.37 | SC130 | Valacyclovir (hydrochloride) | 360.80 |
| SC5 | Acyclovir | 225.20 | SC68 | Isochlorogenic acid A | 516.45 | SC131 | Guanosine | 283.24 |
| SC6 | Oxymatrine | 264.36 | SC69 | Daidzin | 416.38 | SC132 | Sophocarpine (monohydrate) | 264.36 |
| SC7 | Catalpol | 362.33 | SC70 | 2-Phenylethanol | 122.16 | SC133 | Probucol | 516.84 |
| SC8 | Schisandrin C | 384.42 | SC71 | PCL 016 | 123.11 | SC134 | Kaempferide | 300.26 |
| SC9 | Cepharanthine | 606.71 | SC72 | Pyridoxal phosphate | 247.14 | SC135 | Genkwanin | 284.26 |
| SC10 | Oleanonic acid | 454.68 | SC73 | Dipotassium glycyrrhizinate | 861.02 | SC136 | Deapioplatycodin D | 1093.21 |
| SC11 | N6-Methyladenosine | 281.27 | SC74 | Daphnoretin | 352.29 | SC137 | Acetylcysteine | 163.19 |
| SC12 | α-Vitamin E | 430.71 | SC75 | Coptisine (chloride) | 355.77 | SC138 | Chelidonine | 353.37 |
| SC13 | Gentiopicroside | 356.32 | SC76 | Methyl gallate | 184.15 | SC139 | L-Cycloserine | 102.09 |
| SC14 | Shikonin | 288.30 | SC77 | Scutellarin | 462.36 | SC140 | Coumarin | 146.14 |
| SC15 | Valpromide | 143.23 | SC78 | Picroside II | 512.46 | SC141 | 20(R)-Ginsenoside Rh2 | 622.87 |
| SC16 | Rimantadine (hydrochloride) | 215.76 | SC79 | Maslinic acid | 472.70 | SC142 | Mecarbinate | 233.26 |
| SC17 | Impulsin | 299.49 | SC80 | Schisandrin A | 416.51 | SC143 | Azulene | 128.17 |
| SC18 | Brefeldin A | 280.36 | SC81 | Glycitin | 446.40 | SC144 | 3,4'-Dihydroxyflavone | 254.24 |
| SC19 | Cynarin | 516.45 | SC82 | Amentoflavone | 538.46 | SC145 | L-Chicoric Acid | 474.37 |
| SC20 | 2-Deoxy-D-glucose | 164.16 | SC83 | Ivermectin | 875.09 | SC146 | Danthron | 240.21 |
| SC21 | (-)-α-Pinene | 136.23 | SC84 | Aucubin | 346.33 | SC147 | Doxorubicin (hydrochloride) | 579.98 |
| SC22 | Naringenin | 272.25 | SC85 | Corilagin | 634.45 | SC148 | Gomisin G | 536.57 |
| SC23 | Hinokitiol | 164.20 | SC86 | Vidarabine | 267.24 | SC149 | Pentoxifylline | 278.31 |
| SC24 | Dihydrotanshinone I | 278.30 | SC87 | Trigonelline chloride | 173.60 | SC150 | Andrographolide | 350.45 |
| SC25 | Thiamine hydrochloride | 337.27 | SC88 | Harringtonine | 531.59 | SC151 | Mycophenolic acid | 320.34 |
| SC26 | Osthole | 244.29 | SC89 | Oleanolic Acid | 456.70 | SC152 | Sennoside A | 862.74 |
| SC27 | D-Pinitol | 194.18 | SC90 | Octyl gallate | 282.33 | SC153 | Punicalin | 782.53 |
| SC28 | Angelicin | 186.16 | SC91 | Hyperoside | 464.38 | SC154 | Punicalagin | 1084.72 |
| SC29 | N-Acetylneuraminic acid | 309.27 | SC92 | Limonin | 470.51 | SC155 | Verbascoside | 624.59 |
| SC30 | Baicalin | 446.36 | SC93 | Epigoitrin | 129.18 | SC156 | Bergenin | 328.27 |
| SC31 | Ginsenoside Rb1 | 1109.29 | SC94 | Ginsenoside Rb2 | 1079.27 | SC157 | Chlorogenic acid | 354.31 |
| SC32 | Oglufanide | 333.34 | SC95 | Rutin | 610.52 | SC158 | Tunicamycin | 844.94 (n=10) |
| SC33 | Ancitabine (hydrochloride) | 261.66 | SC96 | Aloperine | 232.36 | SC159 | L-Lysine hydrochloride | 182.65 |
| SC34 | Camptothecin | 348.35 | SC97 | Pentagalloylglucose | 940.68 | SC160 | L-Norleucine | 131.17 |
| SC35 | Arctigenin | 372.41 | SC98 | 4-Hydroxyacetophenone | 136.15 | SC161 | Hernandezine | 652.77 |
| SC36 | Lanatoside C | 985.12 | SC99 | Lycorine (hydrochloride) | 323.77 | SC162 | Hanfangichin B (impurified) | 608.74 |
| SC37 | Cephalotaxlen | 315.36 | SC100 | Tizoxanide | 265.25 | SC163 | Ouabain | 584.67 |
| SC38 | Curcumin | 368.38 | SC101 | Valproic acid (sodium salt) | 166.19 | SC164 | N,N'-(hexane-1,6-diyl)bis(2-(2,7-bis(2-(diethylamino)ethoxy)-9H-fluoren-9-ylidene)hydrazine-1-carboxamide) | 1017.38 |
| SC39 | Camphor | 152.23 | SC102 | Efavirenz | 315.68 | SC165 | 2-(2,7-bis(2-morpholinoethoxy)-9H-fluoren-9-ylidene)hydrazine-1-carbothioamide | 511.65 |
| SC40 | Glycyrrhizic acid | 822.93 | SC103 | Betulinic acid | 456.70 | SC166 | 4-((3R,5S,8R,9S,10R,11R,13R,14S,17S)-5,11,14-trihydroxy-10-(hydroxymethyl)-13-methyl-3-(((2S,3S,4R,5R,6S)-3,4,5-trihydroxy-6-methyltetrahydro-2H-pyran-2-yl)oxy)hexadecahydro-1H-cyclopenta[a]phenanthren-17-yl)furan-2(5H)-one | 568.67 |
| SC41 | Dehydroandrographolide succinate | 532.58 | SC104 | Trilobatin | 436.41 | SC167 | 5-chloro-N-(2,6-dichloro-4-nitrophenyl)-2-hydroxybenzamide | 361.57 |
| SC42 | Isomangiferin | 422.34 | SC105 | 4,5-Dicaffeoylquinic acid | 516.45 | SC168 | VE607 | 465.46 |
| SC43 | Emodin | 270.24 | SC106 | Scutellarein | 286.24 | SC169 | SSAA09E3 | 327.34 |
| SC44 | Isoliquiritigenin | 256.25 | SC107 | Valproic acid | 144.21 | SC170 | (2R,2'R)-3,3'-(1,4-phenylenebis(oxy))bis(1-(piperidin-1-yl)propan-2-ol) | 392.54 |
| SC45 | Kaempferol | 286.24 | SC108 | Artemisinin | 282.33 | SC171 | 3,3'-(1,3-phenylenebis(oxy))bis(1-(4-methylpiperidin-1-yl)propan-2-ol) | 420.6 |
| SC46 | Psoralen | 186.16 | SC109 | Raltegravir (potassium salt) | 482.51 | SC172 | 2-(1-(5-chlorothiophen-2-yl)ethylidene)hydrazine-1-carbothioamide | 233.74 |
| SC47 | 9-Aminoacridine | 194.23 | SC110 | Mizoribine | 259.22 | SC173 | N-(4-(4-methylpiperazin-1-yl)benzyl)-4-(pyrrolidin-1-ylmethyl)benzamide | 392.55 |
| SC48 | β-Cyclodextrin | 1134.98 | SC111 | Theaflavin | 564.49 | SC174 | YM-201636 | 467.48 |
| SC49 | Sophocarpine | 246.35 | SC112 | Gramine | 174.24 | SC175 | Amiodarone hydrochloride | 681.77 |
| SC50 | Geldanamycin | 560.64 | SC113 | Artesunate | 384.42 | SC176 | APY0201 | 413.48 |
| SC51 | Xanthone | 196.20 | SC114 | Bevirimat | 584.83 | SC177 | Salinomycin sodium salt | 772.98 |
| SC52 | alpha-Mangostin | 410.46 | SC115 | Elemicin | 208.25 | SC178 | Dronedarone Hydrochloride | 593.22 |
| SC53 | Oroxylin A | 284.26 | SC116 | Fangchinoline | 608.72 | SC179 | Verapamil hydrochloride | 491.06 |
| SC54 | Oxyresveratrol | 244.24 | SC117 | Erythromycin Ethylsuccinate | 862.05 | SC180 | Fangchinoline | 608.71 |
| SC55 | Honokiol | 266.33 | SC118 | Isoferulic acid | 194.18 | SC181 | Isofangchinoline | 608.71 |
| SC56 | α-Lipoic Acid | 206.33 | SC119 | Chebulagic acid | 954.66 | SC182 | Tetrandrine | 622.75 |
| SC57 | Lapachol | 242.27 | SC120 | 4'-O-Methylbavachalcone | 338.40 | SC183 | Berbamine hydrochloride | 681.65 |
| SC58 | Adenosine 5'-monophosphate monohydrate | 365.24 | SC121 | Spermine | 202.34 | SC184 | Dauricine | 624.76 |
| SC59 | Phillyrin | 534.55 | SC122 | Pseudolaric Acid B | 432.46 | SC185 | Neferine | 624.77 |
| SC60 | Hypericin | 504.44 | SC123 | Oxytetracycline | 460.43 | SC186 | Liensinine | 610.74 |
| SC61 | Desaminotyrosine | 166.18 | SC124 | Baicalein | 270.24 | SC187 | Daurisoline | 610.74 |
| SC62 | (-)-Epigallocatechin Gallate | 458.37 | SC125 | Catechin | 290.27 | SC188 | Isoliensinine | 610.75 |
| SC63 | Anthraquinone | 208.21 | SC126 | Cytarabine | 243.22 |  |  |  |

**Table S2**

**Supplementary Tab. 2** The inhibitory activity of 19 hit compounds and aloxistatin against SARS-CoV-2 S-G614 and VSV-G pseudoviruses.

| **Compounds** | **293T-ACE2** | **S-G614 Pseudovirus** | **SI (cytotoxicity)** | **VSV-G Pseudovirus** | **SI (specificity)** |
| --- | --- | --- | --- | --- | --- |
|  | **CC_50_(μM)** | **EC_50_(μM)** |  | **EC_50_(μM)** |  |
| SC8 | ＞50 | 14.58 | ＞3.43 | ﹣ | ﹣ |
| SC9 | ＞50 | 0.32 | ＞158.58 | ＞50 | ＞158.58 |
| SC16 | ＞50 | 34.60 | ＞1.45 | **﹣** | **﹣** |
| SC24 | 33.5 | 15.66 | 0.90 | **﹣** | **﹣** |
| SC33 | ＞50 | 10.12 | ＞4.94 | ＞50 | ＞4.94 |
| SC96 | ＞50 | 0.28 | ＞176.87 | 10.84 | 38.34 |
| SC102 | ＞50 | 0.00038 | ＞130890 | 0.000723 | 1.89 |
| SC109 | ＞50 | 0.00020 | ＞247647 | 0.0359 | 177.86 |
| SC161 | ＞50 | 0.11 | ＞448.83 | ＞50 | ＞448.83 |
| SC171 | ＞50 | 3.83 | ＞13.06 | ＞50 | ＞13.06 |
| SC180 | ＞50 | 0.94 | ＞53.39 | 4.39 | 4.69 |
| SC181 | ＞50 | 1.11 | ＞45.00 | 3.95 | 3.56 |
| SC182 | ＞50 | 0.90 | ＞55.80 | 27.87 | 31.10 |
| SC183 | ＞50 | 1.28 | ＞38.94 | ＞50 | ＞38.94 |
| SC184 | ＞50 | 1.43 | ＞34.99 | ＞50 | ＞34.99 |
| SC185 | ＞50 | 0.95 | ＞52.87 | ＞50 | ＞52.87 |
| SC186 | ＞50 | 1.00 | ＞49.80 | ＞50 | ＞49.80 |
| SC187 | ＞50 | 2.48 | ＞20.13 | ＞50 | ＞20.13 |
| SC188 | ＞50 | 1.29 | ＞38.88 | 7.02 | 5.45 |
| Aloxistatin | ＞50 | 0.923 | ＞54.17 | ＞50 | ＞54.17 |
| The data represent results from three separate experiments | | | | | |
